# GreenLeafVI: A FIJI plugin for high-throughput analysis of leaf chlorophyll content

**DOI:** 10.1101/2025.07.24.666635

**Authors:** Thalia Luden, Jelmer van Lieshout, Sarah L. Mehrem, Basten L. Snoek, Joost Willemse, Remko Offringa

## Abstract

Chlorophyll breakdown is a central process during plant senescence or stress responses and leaf chlorophyll content is therefore a strong predictor of plant health. Chlorophyll quantification can be done in several ways, most of which are time-consuming or require specialized equipment. A simple alternative to these methods is the use of image-based chlorophyll estimation, which uses the color values in RGB images to calculate colorimetric visual indexes as a measure for the leaf chlorophyll content. Image-based chlorophyll measurement is non-destructive and, apart from a digital camera, requires no specialized equipment. Here, we developed the ImageJ plugin GreenLeafVI that facilitates high-throughput image analysis for measuring leaf chlorophyll content. Our plugin offers the option to white-balance images to decrease variation between images and has an optional background removal step. We show that this method can reliably quantify leaf chlorophyll content in a variety of plant species. In addition, we show that image-based chlorophyll quantification can replicate GWAS results based on traditional chlorophyll extraction methods, showing that this method is highly accurate.

## Introduction

Plant health can in many cases be deduced from the leaf chlorophyll (Chl) content, as several stressing factors can affect the photosynthetic capacity of plants. The Chl content can be reduced in response to stresses, such as drought, high salt, nutrient deficiency or pathogen infection, or as a result of programmed plant senescence (Hörtensteiner and Kräutler, 2011). Quantification of Chl is therefore an important tool to monitor plant health in a variety of conditions (Kalaji et al., 2017; Wang et al., 2022). Various methods for measuring Chl content are available, such as solvent-based Chl isolation like acetone extraction followed by photospectrometry at 645 nm and 663 nm (Arnon, 1949), Soil Plant Analysis Development (SPAD) measurement (Konica Minolta Optics, 2009; Markwell et al., 1995; Yadava, 1986), Chl fluorescence measurement (Legendre et al., 2021; Murchie and Lawson, 2013), hyperspectral imaging (Taha et al., 2024; H. Zhang et al., 2022) and image-based colorimetric visual index (CVI) estimations of Chl (Ali et al., 2012; Bresson et al., 2018; Guendouz et al., 2021; Guo et al., 2020a; Taha et al., 2024). The acetone-based Chl extraction method directly measures Chl content, and is one of the most accurate ways to quantify Chl. However, this method is labor-intensive and destructive, and requires specialized equipment to measure the OD645 and OD663. Due to the destructive nature of this method, it is also poorly suited to monitor Chl content over a plant’s life time or under changing conditions. Chl measurements with a SPAD meter are non-destructive, but SPAD measurements remain labor-intensive, as each measurement needs to be recorded manually, and SPAD values are strongly affected by the position on the leaf where the measurements are made. In addition, the SPAD meter can only be used on leaves above a certain size, which excludes the possibility to measure young or small leaves, or plants grown in tissue culture that need to remain in sterile conditions. Chl fluorescence measurements and hyperspectral imaging methods are non-destructive and can yield useful information, but require specialized equipment that can be expensive and are therefore not universally available to researchers. In addition, measuring Chl fluorescence makes use of the photo-saturation of Chl followed by a cool-down period, which makes this type of measurement relatively time-consuming. In contrast, image-based Chl quantification methods require only the use of a digital camera and uniform lighting, are non-destructive and are highly adaptable to different environments and experimental setups. Because no specialized equipment is required, image-based Chl estimation can readily be applied in any environment and is almost universally available to all researchers. Moreover, digital images can be easily processed in batch, allowing high-throughput monitoring of Chl content in a variety of experimental setups (Fernando Sánchez-Sastre et al., 2020; Guendouz et al., 2021; Guo et al., 2020b; Özreçberoğlu and Kahramanoğlu, 2020; Prakash Yadav et al., 2010; Taha et al., 2024; H. Zhang et al., 2022; Zhang et al., 2018).

Image-based Chl estimation relies on the relative pixel intensity in the Red, Green, and Blue channels of RGB images, from which Chl content can be deduced by using different mathematical models (Guo et al., 2020b; Kawashima and Nakatani, 1998; Taha et al., 2024; Woebbecke et al., 1995). Alternatively, values in the hue saturation value color-space (HSV) can be measured (Bresson et al., 2018; Sass et al., 2012). Various studies have shown the usefulness of RGB image analysis and their derived CVIs in estimating Chl content in different species, and strong correlations with total Chl content have been shown for many plant species. For example, Kawashima and Nakatani (1998) showed that the Chl content of wheat and rye leaves could be measured by using a video camera, and showed that the best predictor of Chl content was the normalized difference between the Red and Blue values (R-B)/(R+B), henceforth referred to as the Kawashima index. Other well-performing models that were tested by Kawashima and Nakatani include the normalized Red, Green, and Blue indexes (R/(R+G+B), G/(R+G+B), B/(R+G+B)), and the difference between red and blue divided by the sum of all channels ((R-B)/(R+G+B)). Ali et al. (2012) showed that the Kawashima index correlated strongly with SPAD-502 readings in tomato, and Taha et al. (2024) showed the same in hydroponically-grown lettuce. Similarly, Fernando Sánchez-Sastre et al. (2020) showed that the highest-scoring models tested by Kawashima and Nakatani (1998) also showed a strong correlation with Chl content in sugar beet leaves. On the other hand, Ibrahim et al. (2021) and Ali et al. (2012) found that the Green: Red ratio (G:R) correlated to Chl content in lettuce more strongly than the Kawashima index. Other researchers have used custom models to link the RGB color values to Chl content. For example, Prakash Yadav et al. (2010) showed that Chl content could be accurately measured in micro-propagated potato by using RGB images as well as by the hue, saturation, and intensity values derived from these images, and a similar approach was used in Sorghum by H. Zhang et al. (2022). Other applications of the RGB color space have also been applied, for example by Govindasamy et al. (2017) who used RGB images to monitor the symbiosis efficiency between rhizobia and soybean plants, and Bu et al. (2024) who used RGB parameters to measure soybean pod freshness. Han et al. (2021) used aerial images to monitor the growth of *Hibiscus cannabinus* based on RGB-derived parameters and Zhang et al., (2018) used a similar approach for maize. Finally, Barraza-Moraga et al. (2022) applied RGB analysis to satellite images in order to estimate the Chl-A content of algae in lake Lanalhue in Chile. Overall, these examples show that RGB image analysis can successfully be applied in plant research and is a feasible method of Chl measurement.

Digital phenotyping relies on careful image acquisition and processing. In recent years, several tools have been developed to determine leaf Chl content from digital images, such as the python-based program plantCV (Casto et al., 2022; Gehan et al., 2017) or ImageJ plugins such as the one developed by (Liang et al., 2017). In other cases, researchers made use of custom-made analysis tools such as in MatLab (Ali et al., 2012; Perez-Patricio et al., 2018; Taha et al., 2024). Despite their usefulness, these tools present certain drawbacks for researchers wishing to use RGB images for Chl estimation. PlantCV, while highly customizable and suitable for high-throughput analyses, is based on the Python language and requires a degree of familiarity with this programming language before it can be applied. On the other hand, many biologists are already familiar with the user interface of Fiji/ImageJ, which can be used without prior knowledge of programming languages due to its graphical user interface (Schindelin et al., 2012; Schneider et al., 2012). However, the existing tools available for leaf image analysis in ImageJ either do not measure leaf Chl content (e.g. LeafJ by (Maloof et al., 2013)), or require several manual image calibration steps (Liang et al., 2017), making them unsuitable for high-throughput analysis of Chl. To our knowledge, there is no tool offering both high-throughput analysis of RGB images that offers a direct estimation of Chl content and that does not require programming skills. We therefore set out to develop an ImageJ plugin that reliably estimates leaf Chl content based on digital images in a high-throughput manner for various applications.

The open-source software FIJI is a distribution of the image analysis program ImageJ, which is a commonly used and highly customizable tool for image analysis in the life sciences (Schindelin et al., 2012). FIJI offers possibilities to automate analysis steps via the use of macros or custom-made plugins. In order to enhance the throughput of RGB image analysis, we developed the FIJI plugin called Green Leaf Visual Index (GreenLeafVI) that can perform multiple steps of RGB analysis in batch mode for a large number of images, allowing for high-throughput analysis. GreenLeafVI offers the option to normalize image brightness, segment images to reduce background noise and calculate the RGB values of multiple objects per image along with various methods of leaf Chl content estimation. The output data is stored in a tidyR-compatible format that can readily be used for further statistical analysis in R or other statistics software. We show that GreenLeafVI can be applied for measuring Chl content of different plant species and can accurately reproduce GWAS results obtained by traditional Chl measurement methods in lettuce, validating its use for high-throughput phenotyping experiments.

## Methods

### Plant materials and growth conditions

*Arabidopsis thaliana* (Arabidopsis) ecotype Col-0 seeds were sown on humid soil (90% turf, 10% sand) and stratified at 4°C for three days, and subsequently transferred to a growth chamber with long day conditions (16h light / 8h dark) at 21°C and 65% relative humidity. Seedlings were repotted to individual pots after seven days, and the fifth leaf of each plant was harvested at 23 days after the end of stratification. Leaves incubated for zero to five days in the dark were used for senescence measurements to represent various stages of leaf senescence.

For testing the correlation between colorimetric measurements and Chl content, *Lactuca sativa* (lettuce) cv. Cobham Green seeds were sterilized in 50% chlorix bleach and 50% MilliQ solution solution and stratified in distilled water for four days at 4°C. Subsequently, seeds were sown on moist soil (90% turf, 10% sand) and transferred to a growth chamber with long day conditions (16h light / 8h dark) at 21°C and 70% relative humidity. The fourth leaf was harvested ten days after emergence, and 25 mm Ø leaf disks were taken and incubated on demineralized water with 1% agarose in the dark for up to five days. Leaves subjected to different dark incubation periods were used for senescence measurements to represent a range of senescence stages. For the GWAS experiment, we selected a total of 184 *Lactuca sativa* accessions as described by R F Dijkhuizen et al.(2024). Seeds were treated as described above, and leaf disks of the fourth leaf from three plants per cultivar were taken at 10 days after leaf emergence and used directly for imaging and Chl extraction.

*Nicotiana benthamiana* (tobacco) seeds were sown on moist soil (90% turf, 10% sand) and germinated in a growth chamber with long day conditions (16h light / 8h dark) at 21°C and 70% relative humidity. After seven days, seedlings were repotted to individual pots. After three weeks, mature leaves were detached and incubated in large petri dishes on demineralized water with 0,5% agarose in the dark for up to seven days.

*Solanum lycopersicum* (tomato) cv. Moneymaker seeds were sterilized in bleach and germinated on solid MS medium, and transferred to soil after three weeks. Plants were grown in long days conditions (16h light / 8h dark) at a 24°C day / 18°C night temperature regime. Leaves were harvested from flowering plants and incubated in large petri dishes on demineralized water with 0.5% agarose in the dark for up to seven days, with harvesting points between zero to seven days of dark incubation.

### Senescence induction, imaging, and Chl isolation

Arabidopsis leaves were floated on 5 mM MES buffer (pH 5.6) in 5 cm petri dishes sealed with parafilm, wrapped in aluminum foil, and kept at 21°C for up to five days. Lettuce, tomato, and tobacco leaves were harvested at various ages and placed in petri dishes with demineralized water with 0.5% agarose, sealed with parafilm, wrapped in aluminum foil, and stored at 21°C for up to seven days. Leaves were imaged at different time points between zero and seven days of dark incubation and leaf disks were collected for Chl extraction after imaging. All images were taken with a Nikon DC3000 DSLR camera under uniform white light. Plant leaves were placed on a homogenous white or black background along with a white square that served as a reference for image brightness.

Chl extraction was performed according to the protocol of Arnon (1949). Briefly, 5 mm diameter round leaf disks were taken after imaging (2 per Arabidopsis leaf and 3 per leaf for other species) and placed in a 96-well deep-well plate along with a metal bead and stored at -80°C until extraction. For extraction, the leaf tissue was pulverized and resuspended in 200 μL 25 mM sodium phosphate buffer (pH 7) and 800 μL 80% (v/v) acetone. Samples were then incubated at room temperature in the dark for 1 hour with gentle shaking and centrifuged for 10 minutes at 3000 *g*. 200 μL of the supernatant was then transferred to a 96-well transparent-bottom plate and the absorption (D) at 645nm and 663 nm was measured with a Spark 10M microplatereader (TECAN Group AG, Männedorf, Switzerland). The total Chl content in mg·L^-1^ was then calculated with the following formula: Chl_Total_ = 20,2·D_645_ + 8,02·D_663_ as described by Arnon (1949). We decided to measure the Chl content in mg·cm^-2^ rather than mg·g^-1^ fresh weight to minimize variation between measurements because we observed that leaves wilted after several days of dark incubation, reducing the fresh weight whereas leaf area was less affected by the senescence process. To calculate the Chl in mg·cm^-2^, the amount of Chl in one mL (the extraction volume for each sample) was divided over the total area of leaf tissue used for extraction.

### Image analysis

All images were analyzed using the FIJI open source release of ImageJ2 (Schindelin et al., 2012), with the custom GreenLeafVI plugin that can perform white balancing, automatic selection of leaves and removal of background, and RGB pixel intensity measurements semi-automatically. The plugin is described in detail in Supplementary protocol 1, and Figure 1 shows a schematic overview of the steps performed by the plugin.

**Figure 1:**
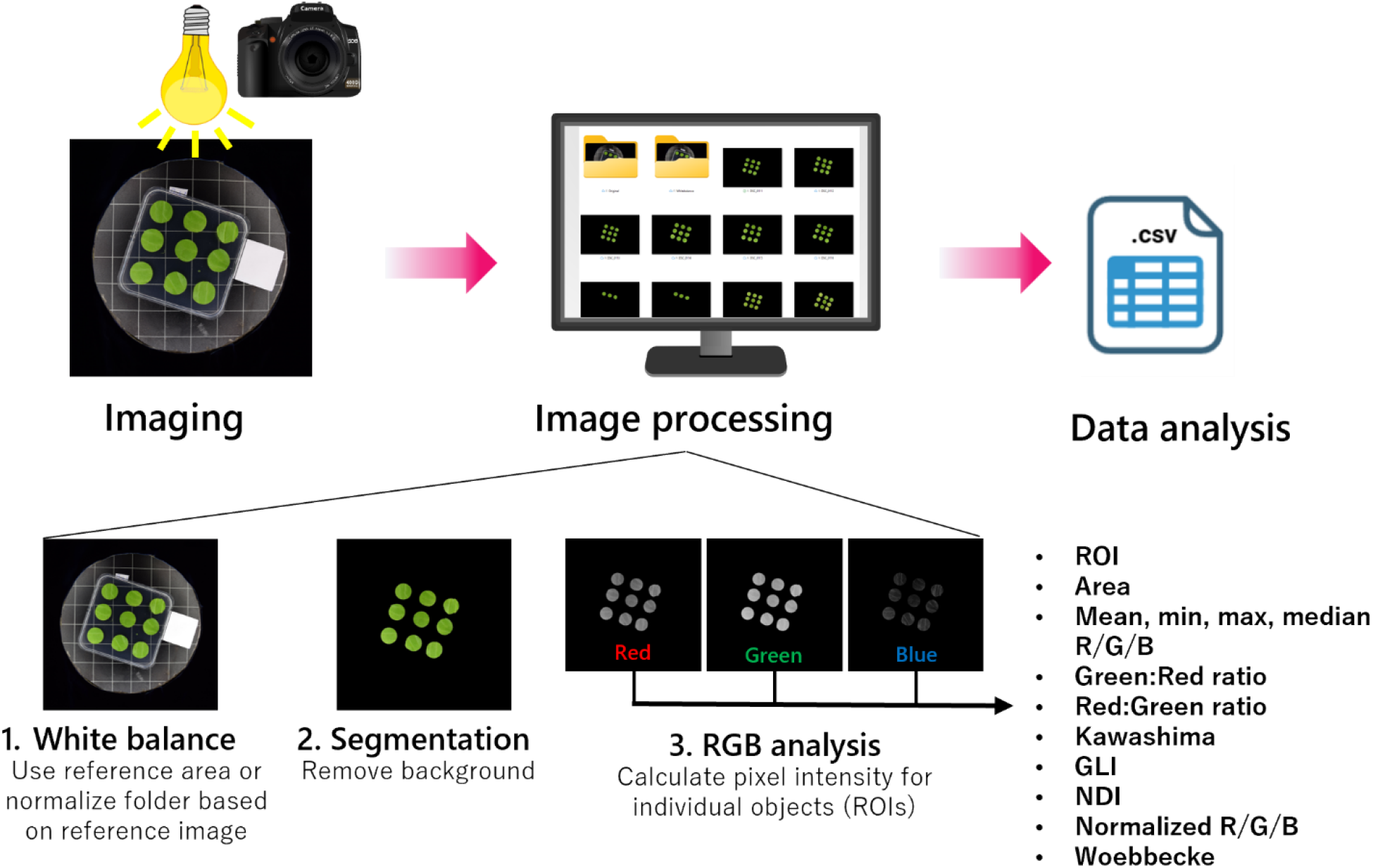
Workflow for high-throughput Chl measurement with GreenLeafVI. Images are taken under homogenous light with a white reference area. The images are then batch-processed for 1) white-balancing based on the white reference area; 2) segmentation to remove the background; and 3) measurement of pixel intensity in the Red, Green, and Blue channels. The results of the RGB analysis are stored in a results.csv file, which, in addition to the RGB values, also contains the region of interest (ROI), the area of the objects, and several colorimetric visual indices (CVIs).

#### White balancing

In order to calibrate the image brightness and reduce inter-image variation, we normalized the pixel intensity to the white reference area included in each image (Figure 1). This was done by splitting the image into Red, Green, and Blue channels and automatic measurement of the mean pixel intensity in the white reference area. Next, the pixel adjustment factor for each of the channels was calculated by dividing the maximum brightness (255 for 8-bit images) by the mean pixel intensity of the white area (adjustment factor = 255/mean). The white reference was then set to an intensity of 255 in all three channels, and the pixels outside of the reference area were normalized based on the adjustment factor for each channel, and re-stacked into an RGB image with a “_whitebalanced” suffix for further analysis.

#### Segmentation of images to reduce background

In order to exclude measurements of non-leaf objects or areas in the background of the image, we included an optional segmentation step in the GreenLeafVI plugin (Figure 1). The segmentation step extracts leaf-like objects from the background by creating a mask covering the leaves based on minimum and maximum Hue Saturation Value (HSV) and minimum area parameters, and removes the background by setting the pixel intensity outside of masked areas to 0. The HSV values used to distinguish leaves from the background were calibrated manually for each species. The images were then saved extended by a “_segmented” suffix for further analysis. We choose HSV as a color model for segmentation as it not only filters for color (Hue), but also incorporates intensity (Saturation) and lightness (Value), improving object from background segmentation.

#### RGB analysis and color-based methods of Chl estimation

To measure the R, G, and B pixel intensities of individual objects in an image, a mask selecting individual objects was made like in the segmentation step. The original image was then split into Red, Green, and Blue channels and the minimum, maximum, median, and mean pixel intensity was measured in each object of the mask covering the leaves (Figure 1). The mean values of each leaf measured in the Red, Green, and Blue channels were used to calculate the different visual indexes: Green:Red ratio (GR_ratio), Red:Green ratio (RG_ratio), Kawashima index (Kawashima), Green Leaf Index (GLI), Normalized Difference Index (NDI), normalized Red (Red_norm), normalized Green (Green_norm), normalized Blue (Blue_norm), and the Woebbecke index (Woebbecke).

### Data analysis

#### Correlation analyses

Data generated by the ImageJ plugin and from spectrophotometry results were analyzed in R (R Core Team, 2023). Pearson’s correlation analysis between the different colorimetric indexes and the total Chl content as mg Chl·cm^-2^ was performed with the R package psych (William Revelle, 2023), and plots were generated by using the ggplot2 (Wickham, 2016) and ggpubr packages (Kassambara, 2023).

#### GWAS analysis

SNP data was obtained from Dijkhuizen et al. (2024). Kinship was computed as the covariance matrix of SNPs using the cov() function in R. SNPs were filtered for a MAF > 0.05. For GWAS we used R version 4.2.2 and the lme4QTL package (Ziyatdinov et al., 2018). GWAS was performed on the filtered SNP set using a linear mixed model (relmatLmer and matlm), with the kinship matrix included as a random effect. The Bonferroni method was used to correct for multiple testing, with the Bonferroni-corrected significance threshold being - log10(0.05/2,485,803) > 7.69. Manhattan plots were created using ggplot (Wickham, 2016)(Wickham, 2016). Quantile-Quantile (Q-Q) plots were created using the ggfastman R-package (Tremmel, 2021). The genome annotation and gene function prediction of the Salinas reference genome V8 were used for annotation of significantly associated loci (Reyes-Chin-Wo et al., 2017).

## Results and discussion

### Normalized Red and Green:Red ratio are accurate predictors of the Chl content in leaves

We designed the Green Leaf Visual Index (GreenLeafVI) plugin as a tool to easily phenotype the Chl content in different plant species. We therefore ran the three steps of the GreenLeafVI plugin (white balancing, segmentation, and RGB analysis) on four different plant species: Arabidopsis, tobacco, tomato and lettuce. In addition, we measured the Chl content in extracts from each imaged leaf in order to test how well the different CVIs measured by the GreenLeafVI plugin correlated to the actual Chl content. To assess which CVI reflects actual Chl content most accurately, we performed linear regression analysis with a number of previously described CVIs and the actual Chl content in mg·cm^-2^ for the four plant species. Our data show that several CVIs show a strong correlation with the total Chl content for these different plant species (Figures 2A-D). Overall, the CVI-based prediction of Chl was most accurate for lettuce, where the Green: Red ratio (G:R) and the Normalized Difference Index (NDI; (Rn-Gn)/(Rn+Gn+0.01)) were the most robust predictors of Chl content followed by the normalized Red value (Rn; R/(R+G+B)) (Figure 2D, Table 1). For Arabidopsis, tobacco, and tomato the normalized Red value best reflected the Chl content, although we found that Chl also strongly correlated with the NDI and G:R (Figure 2A-C, Table 1). Interestingly, we observed marked differences in CVI accuracy between species, with lettuce showing the best overall correlation between the actual Chl content and most CVIs, and tobacco the weakest, suggesting that CVIs are more suitable for comparing Chl levels in some species compared to others. Previous research has shown that G:R is a good indicator of Chl in lettuce (Taha et al., 2024) and that Rn shows the best consistency between species (Ali et al., 2012). Taken together, our data show that overall, Rn, G:R, and NDI are the best predictors of actual Chl content and can be reliably used to compare Chl levels in different samples.

**Table 1:** Correlations of various CVIs with the chlorophyll content in mg Chl·cm^−2^ (linear regression). In the formula column, R, G, and B stand for average Red, Green, and Blue pixel intensity values. Curves were fitted with a linear model fitting *y* ∼ *x*, with y representing mg Chl/cm2 and x the CVIs in all cases except for the Woebbecke Index (*), where the curve was fit to a model of 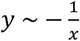.

| Index | Formula | Arabidopsis |  | Tomato |  | Tobacco |  | Lettuce |  |
| --- | --- | --- | --- | --- | --- | --- | --- | --- | --- |
|  |  | R2 | p value | R2 | p value | R2 | p value | R2 | p value |
| Green:Red ratio | G/R | 0.607 | <2.2e-16 | 0.739 | 4.55E-08 | 0.3627 | 6.10E-05 | **0.752** | **1.61E-14** |
| Kawashima Index | (R-B)/(R+B) | 0.205 | 1.33E-13 | 0.551 | 2.00E-05 | -0.027 | 0.7600 | 0.195 | 0.0016 |
| Normalized Red (Rn) | R/(R+G+B) | **0.647** | **<2.2e-16** | **0.775** | **8.62E-09** | **0.590** | **2.72E-08** | 0.673 | 5.92E-12 |
| Normalized Green (Gn) | G/(R+G+B) | 0.343 | <2.2e-16 | 0.481 | 1.04E-04 | 0.204 | 0.0034 | 0.646 | 3.11E-11 |
| Normalized Blue (Bn) | B/(R+G+B) | 0.009 | 0.0755 | 0.092 | 0.0821 | 0.005 | 0.2876 | 0.363 | 9.02E-06 |
| Normalized Difference Index (NDI) | (Rn-Gn)/(Rn+Gn+0.01) | 0.608 | <2.2e-16 | 0.739 | 4.45E-08 | 0.370 | 4.98E-05 | **0.754** | **1.37E-14** |
| Green Leaf Index (GLI) | (2G-R-B)/(2G+R+B) | 0.339 | <2.2e-16 | 0.487 | 9.10E-05 | 0.203 | 0.0034 | 0.649 | 2.60E-11 |
| Woebbecke Index* | (G-B)/(R-G) | 0.638 | <2.2e-16 | 0.822 | 6.603E-10 | 0.513 | 5.49E-07 | **0.768** | **4.03E-15** |

**Figure 2:**
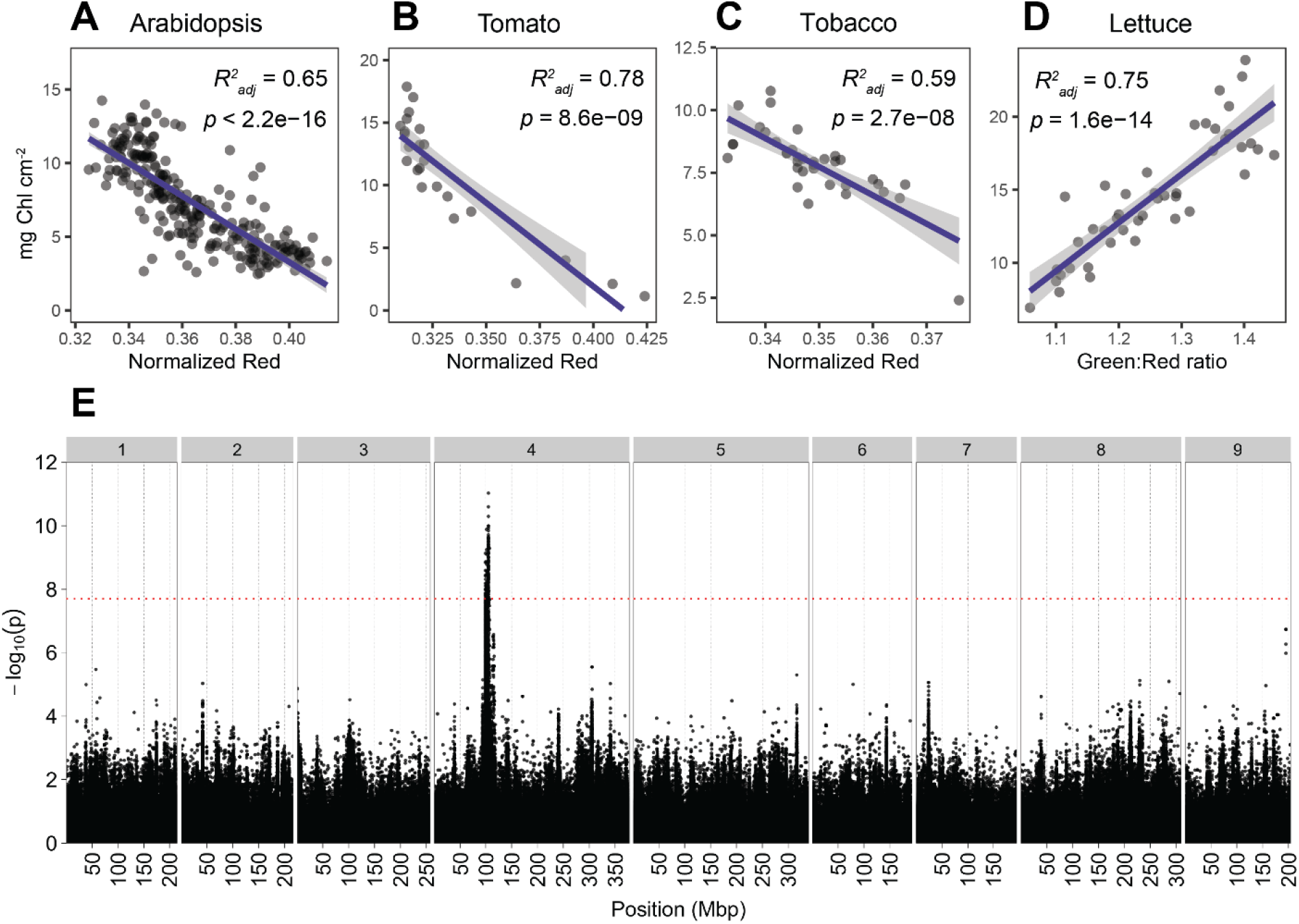
Application of CVIs for measuring Chl content in different species. **A-D**: correlations between total Chl content as mg Chl·cm^-2^ and the best-scoring CVI for each species. **A**: Arabidopsis, **B**: tomato and **C**: tobacco each show the strongest correlation between mg Ch·cm^-2^ and the normalized Red value (Rn). **D**: In lettuce, mg Chl·cm^-2^ correlates best with the Green: Red ratio (G:R). **E**: Manhattan plot of the GWAS on the Green: Red ratio measured on the fourth leaf of 184 lettuce cultivars. Genomic position, indicated in megabasepairs (Mbp), is shown on the x-axis, chromosome numbers are indicated on top. Significance as – log_10_(p) is shown on the y-axis. The Bonferroni threshold of -log_10_(p) > 7.69 is indicated by the red horizontal line.

Surprisingly, the correlation between the Chl content and the Kawashima index was not as strong as shown in some earlier reports e.g. (Kawashima and Nakatani, 1998; Taha et al., 2024), and was outperformed by G:R, NDI, and Rn in all four species. Similar results were obtained by (Guo et al., 2020b), who showed that both the Kawashima and Woebbecke Indexes were outperformed by G:R and GLI when using aerial RGB images to estimate the Chl content in field-grown maize. Ali et al., (2012) showed that the Kawashima index performed well in tomato, but not in lettuce or broccoli, whereas Rn performed well and was most consistent between species, a finding that is reinforced by our results. One possible explanation for the poor performance of the Kawashima index compared to Rn and other indexes could be the omission of G in the Kawashima index formula ((R-B)/(R+B); Table 1), despite the green color of Chl. This corresponds with the fact that each of the best-performing indexes in our setup (Rn, G:R, NDI) included green pixel intensity in their formulas (Table 1), showing that the green value is of importance for Chl estimation. However, greenness alone was not sufficient to accurately predict the Chl content, as the normalized Green value (Gn; G/(R+G+B)) showed only a moderate correlation with the actual Chl content in our experiments (Table 1) as well as in previous studies (Ali et al., 2012). Together with our finding that Rn and G:R are the top performing CVIs, it is clear that the intensity in the green and especially red channels best reflect the actual Chl content, whereas the intensity in the blue channel provides additional, but not indispensable, information.

Upon initial linear regression analysis, the Woebbecke Index appeared to correlate very poorly with the Chl content. While most CVIs showed a linear correlation with the actual Chl content of the type y = x (with x = CVI, y = mg Chl·cm^-2^, Figures S1-7), the Woebbecke Index appeared to form a non-linear, hyperbolic relation with the actual Chl content (Figure S8). A linear model using a regression curve fitting with a formula of the type y = -1/x (with x = Woebbecke Index, y = mg Chl·cm^-2^) showed that the Woebbecke Index actually correlated more strongly than any other CVI with the Chl content in tomato and lettuce, with an *R*^*2*^ of 0.801 in tomato and an *R*^*2*^ of 0.768 in lettuce (Figure S2, Table 1). However, despite the high *R*^*2*^ values implying the Woebbecke Index as the superior CVI, we observed that the Woebbecke index sometimes caused extremely positive or negative values that were not reflected by the other CVIs or actual Chl content, which could complicate comparisons between different measurements. While for lettuce and tobacco the Woebbecke Index generated exclusively negative values in a relatively narrow range, we observed that in senescent leaves of Arabidopsis and tomato the Woebbecke Index formula generated extremely low (<-200) or extremely high (>400) measurements (Figure S8). The Woebbecke Index formula is defined as the difference between the G and B channels divided by the difference between R and G ((G-B)/(R-G); Table 1). Because of this, the Woebbecke Index can create two types of abnormal values: extreme outliers and unexpected positive values.

In cases where the R and G values are similar, such as in senescent leaves, the denominator of this formula approaches 0, resulting in extreme outlier values that may be either positive (when R>G≥B or B>G≥R) or negative (when B>G and R≥G or G>B and G≥R). In addition, more mild but still unexpected positive values can be generated in situations where R>G>B or B>G>R). These unexpected positive values or extreme values complicate the comparison between groups, as the mean value of the measurements in one group will be strongly affected by such values. Therefore, while the Woebbecke Index shows a strong hyperbolic correlation with mg Chl·cm^-2^ and might be suitable to compare greenness among healthy, non-senescent plants where R and G values are sufficiently different, our results show that it is unsuitable for comparing Chl in senescent, yellowing leaves. We therefore advise to use more robust indexes such as Rn, NDI, or G:R.

For this study, we used leaves at various stages of senescence (non-senescent up to completely senesced) for Chl measurements and imaging. By including leaves in a range of senescence stages, we generated a dataset that that is representative for a variety of conditions, where yellowing can represent different types of stress. Previous research such as (Liang et al., 2017; Woebbecke et al., 1995) largely focused on measurement of the Chl content in healthy and/or young plants in order to monitor plant health during early development. Although this has yielded well-performing CVIs, the omission of senescent or otherwise yellow leaves in the index calibration could explain why some indexes (e.g. Kawashima or Woebbecke) perform less well in our conditions compared to previous studies. In addition, most studies have made use of SPAD readings to calibrate or measure the success of a CVI (Ali et al., 2012; Guendouz et al., 2021; Taha et al., 2024; Wang et al., 2022; Yuan et al., 2022), whereas we have directly measured the fluorescence of acetone-extracted Chl. Although the SPAD value and extraction-based Chl quantification methods correlate strongly, with *R*^*2*^ values larger than 0.9 (Castelli et al., 1996; Markwell et al., 1995), the two methods still show slight differences, which could partially explain why some CVIs perform better in our study compared to previous research and vice versa. Despite these differences, our data shows that several visual indexes perform well in predicting Chl content, and that the Rn, G:R, and NDI indexes are all reliable, with slight differences in the top performing CVI for each species.

### Use of GreenLeafVI for GWAS on the Chl content in lettuce

In order to test whether visual indexes are sufficiently accurate to identify phenotypic differences between genotypes, we ran a Genome-Wide Association Study (GWAS) on leaf Chl content in lettuce, the plant species where the use of digital images correlated most strongly with actual Chl content. We used a panel of 184 lettuce cultivars grown in controlled conditions, and harvested the fourth leaf at 10 days after emergence for three plants of each genotype. We then imaged a 30 mm leaf disk of this leaf, ran the GreenLeafVI plugin on the images thus generated, and used the results for subsequent analyses. We used an extensive SNP dataset to run GWASs on G:R (the best-performing CVI for lettuce) and Rn (the overall best performing CVI) as a measure of Chl content. From these GWASs, a significant peak on chromosome 4 was identified (Figure 2E, Figures S9-10), which corresponded to the peak found by L. Zhang et al. (2022) in their GWAS on 125 lettuce genotypes using Chl measurements on leaf extracts. This peak was shown to be associated with the lettuce *Golden-Like (LsGLK)* gene, which is an important regulator of chloroplast development (L. Zhang et al., 2022). The association of the significant peak that was observed in all three GWASs with a gene regulating chloroplast development explains the variation in Chl content that was observed by either direct Chl measurement (L. Zhang et al., 2022) or, in our case, by using digital images as a reference. The similarity in outcomes between the two GWASs clearly shows that our G:R and Rn data can reproduce the association between the Chl content phenotype and the SNP located near the *LsGLK* gene, indicating that G:R is not only an accurate measure of Chl content, but is also a reliable method to identify a phenotypic vs genotypic association.

### Concluding remarks

In addition to their accuracy at reflecting Chl content, digital images can be easily stored and re-examined at a later time point. Other traits besides Chl content (e.g. anthocyanin production, leaf shape and size, etc.) can be deduced from digital images, and such traits may be examined at any moment when such images are stored with proper documentation, whereas this information is lost when using destructive methods of Chl quantification. Thus, image-based Chl measurement offers additional benefits compared to direct Chl measurement. Making digital images and automated image processing is also less labor-intensive and less costly than large-scale Chl extraction. Compared to existing methods to measure RGB values in leaves or plants, the GreenLeafVI plugin presented here is easy to use and can help researchers measure and monitor the Chl content in a variety of species in a non-destructive way. Considering the various benefits of digital image-based Chl measurement, we propose that this method be used in future applications.

## Author contributions

TL, JvL and RO designed the experiments. TL and JvL performed Chl extraction experiments, and JvL performed lettuce phenotyping for GWAS. Correlation analyses were done by TL, and SLM and BS performed GWAS analysis. JvL, TL, and JW wrote the ImageJ macro and java scripts. TL wrote the manuscript with input from all other authors.

## Acknowledgements

We would like to thank Jan Vink, Altay Temel, Ward de Winter and Mariel Lavreijsen for technical support. We also thank BSc/MSc students Marion Larue, Karin Verkerk and Kim Roos for their help in phenotyping.

## Data availability

The data that support the findings of this study are available from the corresponding author upon reasonable request. The GreenLeafVI source code, documentation and further information is available at https://github.com/jelmervanlieshout/GreenLeafVI.

## Supporting information

**Figure S1**: Correlation between Green: Red ratio and Chl content in Arabidopsis, tomato, tobacco, and lettuce.

**Figure S2:** Correlation between the Kawashima Index and Chl content in Arabidopsis, tomato, tobacco, and lettuce.

**Figure S3:** Correlation between the Normalized Red value (Rn) and Chl content in Arabidopsis, tomato, tobacco, and lettuce.

**Figure S4:** Correlation between the Normalized Green value (Gn) and Chl content in Arabidopsis, tomato, tobacco, and lettuce.

**Figure S5:** Correlation between Normalized Blue value (Rn) and Chl content in Arabidopsis, tomato, tobacco, and lettuce.

**Figure S6:** Correlation between the Normalized Difference Index (NDI) and Chl content in Arabidopsis, tomato, tobacco, and lettuce.

**Figure S7:** Correlation between the Green Leaf Index (GLI) and Chl content in Arabidopsis, tomato, tobacco, and lettuce.

**Figure S8:** Correlation between the Woebbecke Index and Chl content in Arabidopsis, tomato, tobacco, and lettuce.

**Figure S9**: Quantile-Quantile plot for Green: Red ratio on the fourth leaf of 184 lettuce cultivars.

**Figure S10**: Manhattan plot of the GWAS on the normalized Red value measured on the fourth leaf of 184 lettuce cultivars.

**Protocol S1**: User guide for GreenLeafVI plugin.

## Supporting information

**Figure S1:**
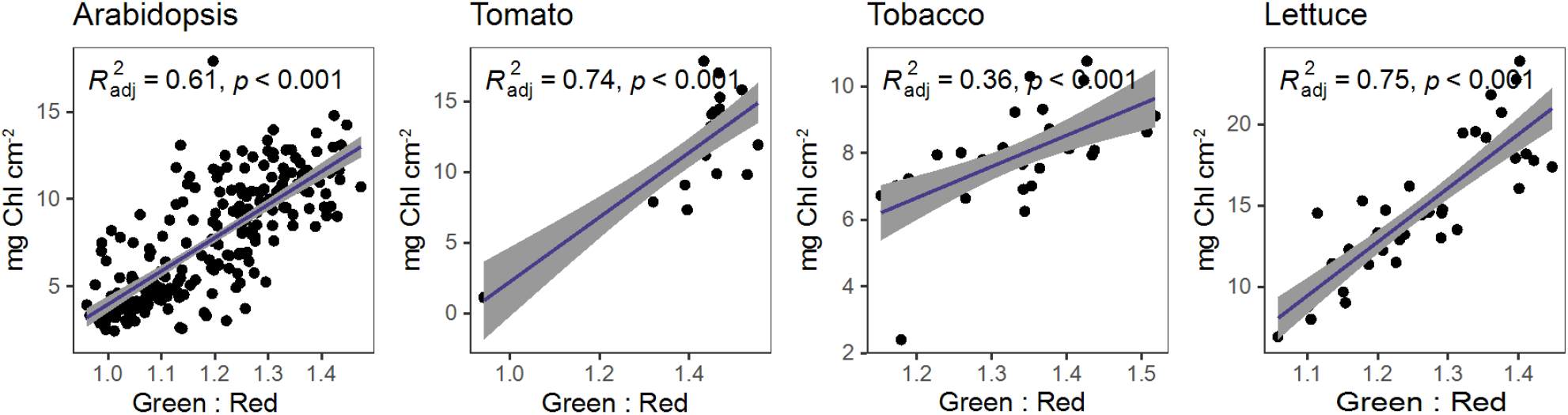
Correlation between Green: Red ratio and Chl content in Arabidopsis, tomato, tobacco, and lettuce. Correlation analysis was done by linear regression using a model where *y* = *x*.

**Figure S2:**
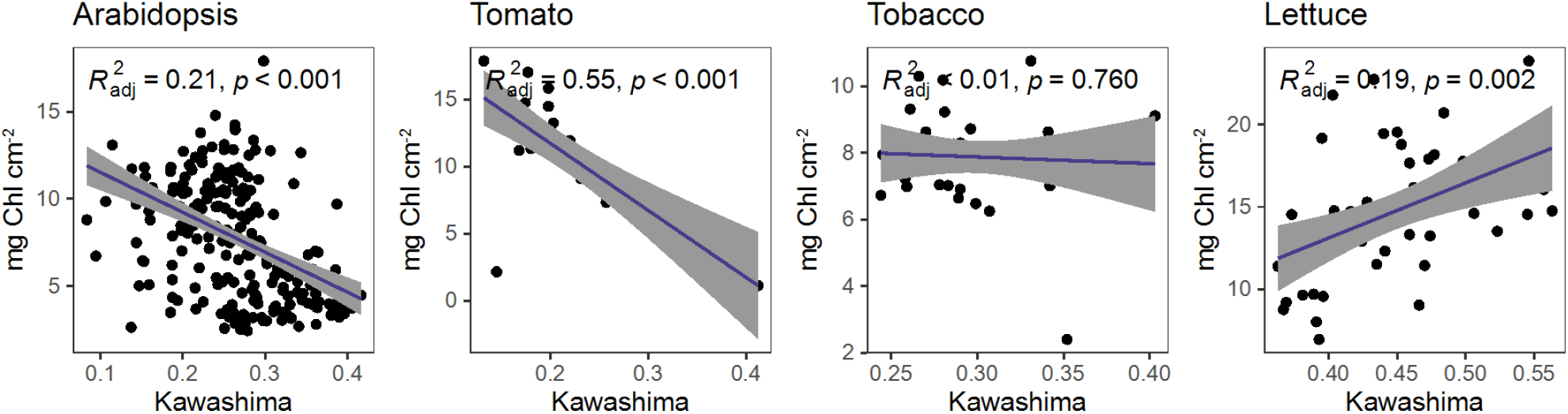
Correlation between the Kawashima Index and Chl content in Arabidopsis, tomato, tobacco, and lettuce. Correlation analysis was done by linear regression using a model where *y* = *x*.

**Figure S3:**
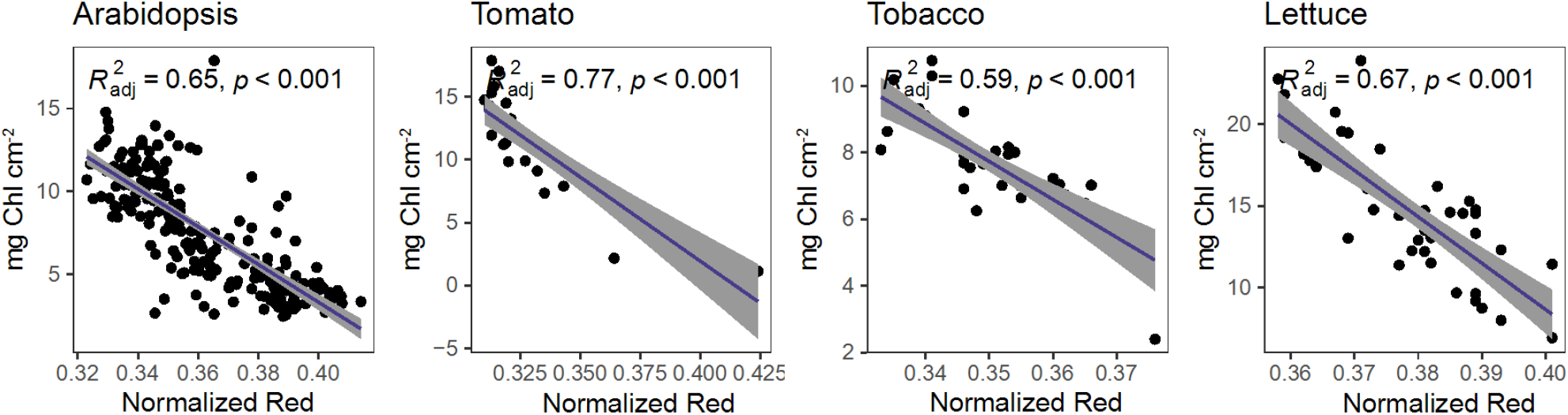
Correlation between the Normalized Red value (Rn) and Chl content in Arabidopsis, tomato, tobacco, and lettuce. Correlation analysis was done by linear regression using a model where *y* = *x*.

**Figure S4:**
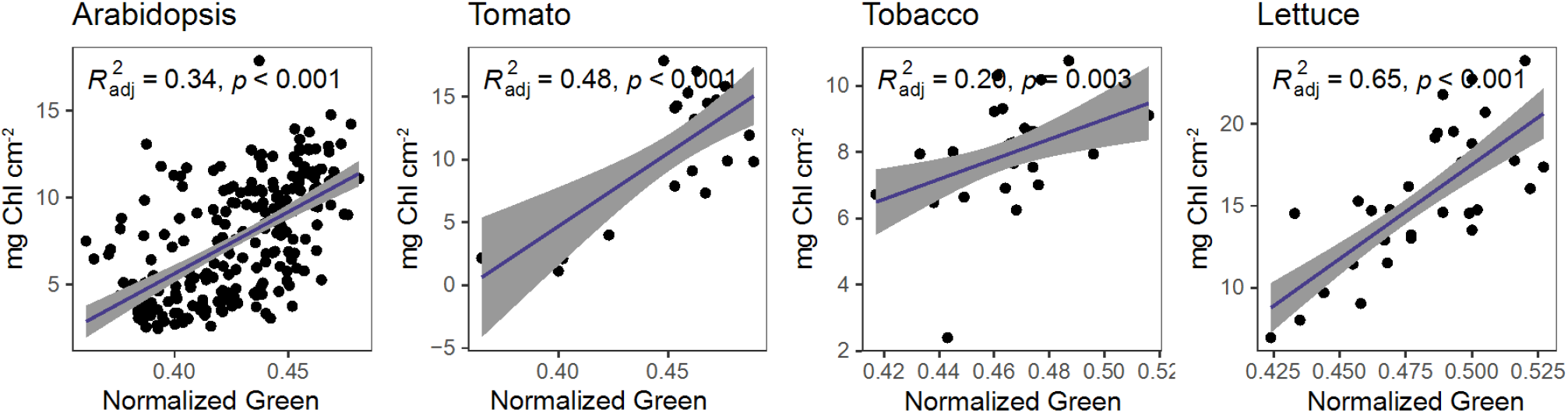
Correlation between the Normalized Green value (Gn) and Chl content in Arabidopsis, tomato, tobacco, and lettuce. Correlation analysis was done by linear regression using a model where *y* = *x*.

**Figure S5:**
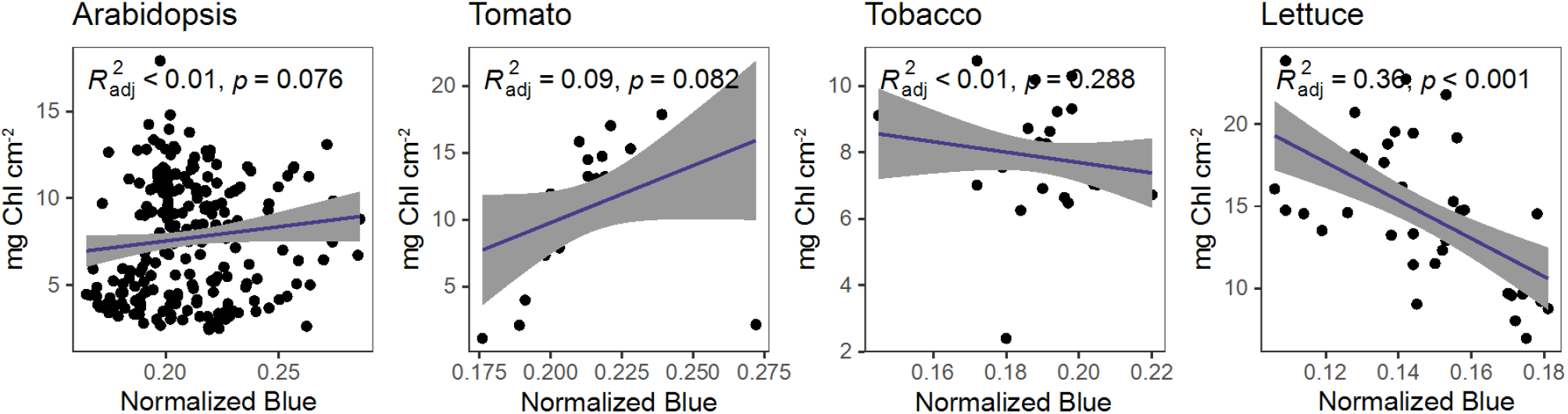
Correlation between Normalized Blue value (Rn) and Chl content in Arabidopsis, tomato, tobacco, and lettuce. Correlation analysis was done by linear regression using a model where *y* = *x*.

**Figure S6:**
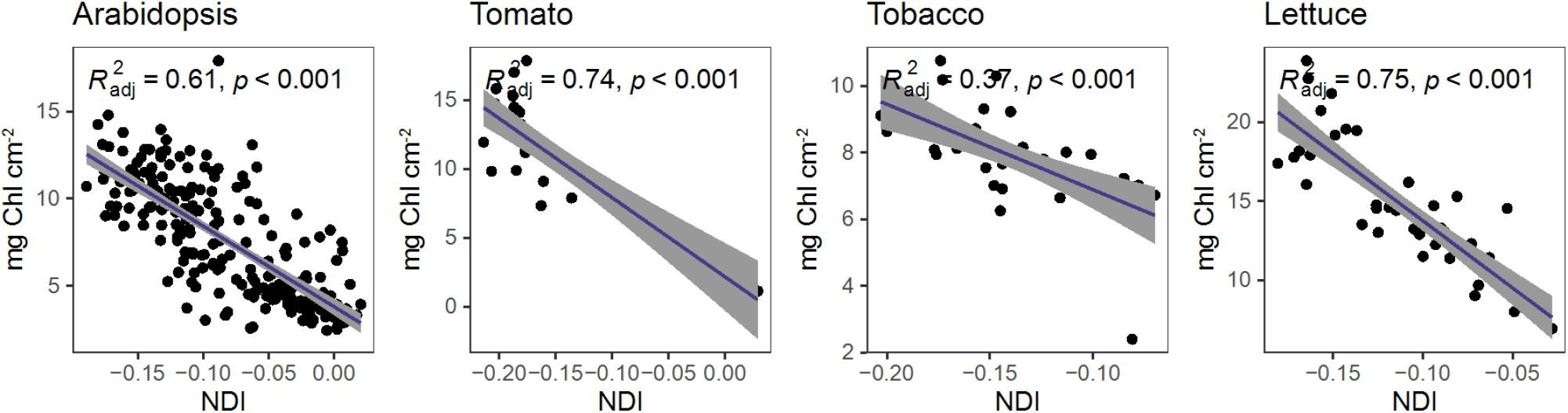
Correlation between the Normalized Difference Index (NDI) and Chl content in Arabidopsis, tomato, tobacco, and lettuce. Correlation analysis was done by linear regression using a model where *y* = *x*.

**Figure S7:**
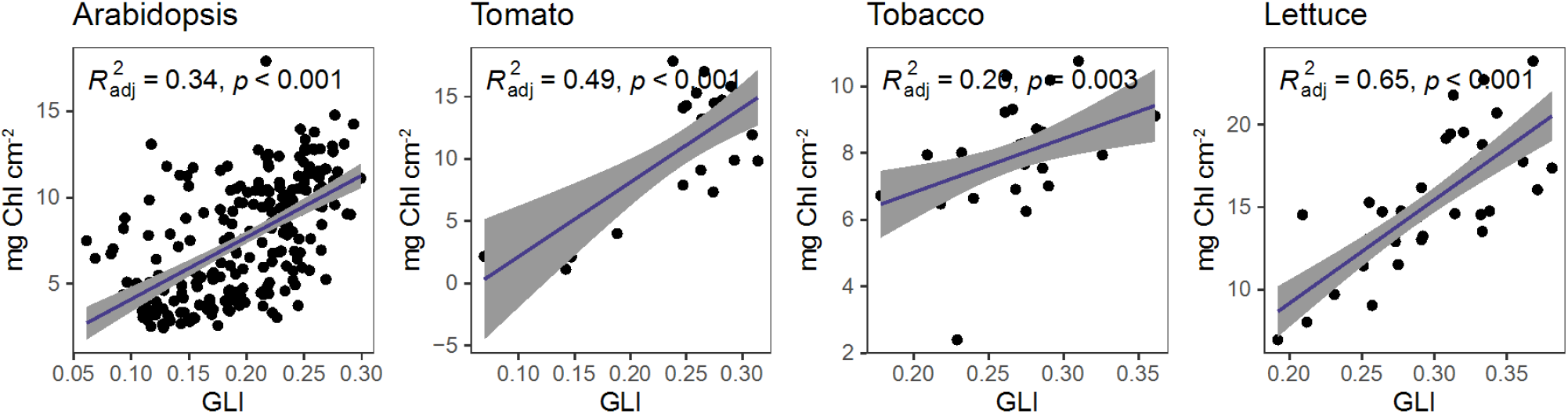
Correlation between the Green Leaf Index (GLI) and Chl content in Arabidopsis, tomato, tobacco, and lettuce. Correlation analysis was done by linear regression using a model where *y* = *x*.

**Figure S8:**
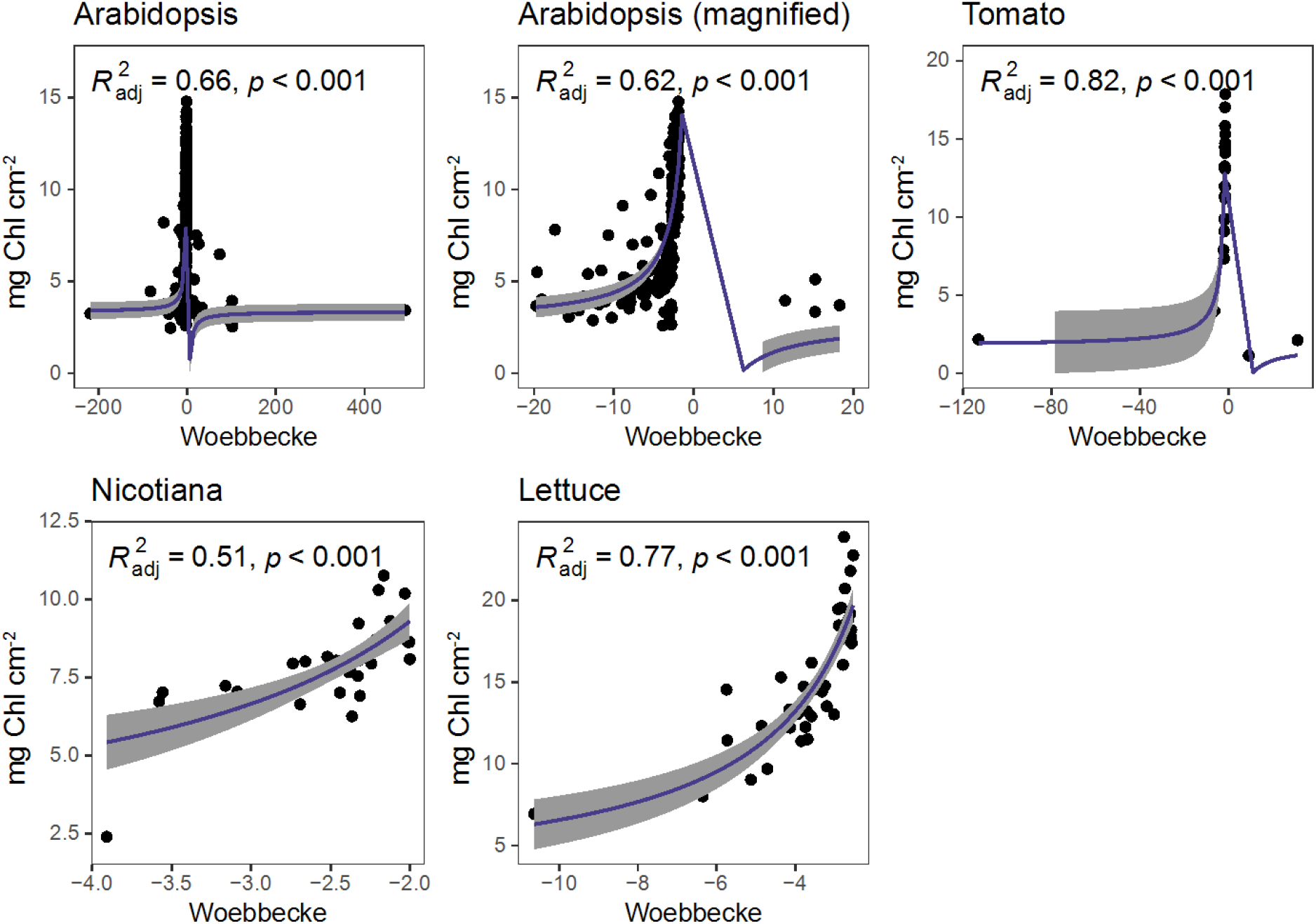
Correlation between the Woebbecke Index and Chl content in Arabidopsis, tomato, tobacco, and lettuce. Correlation analysis was done by linear regression using a model where 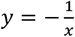.

**Figure S9:**
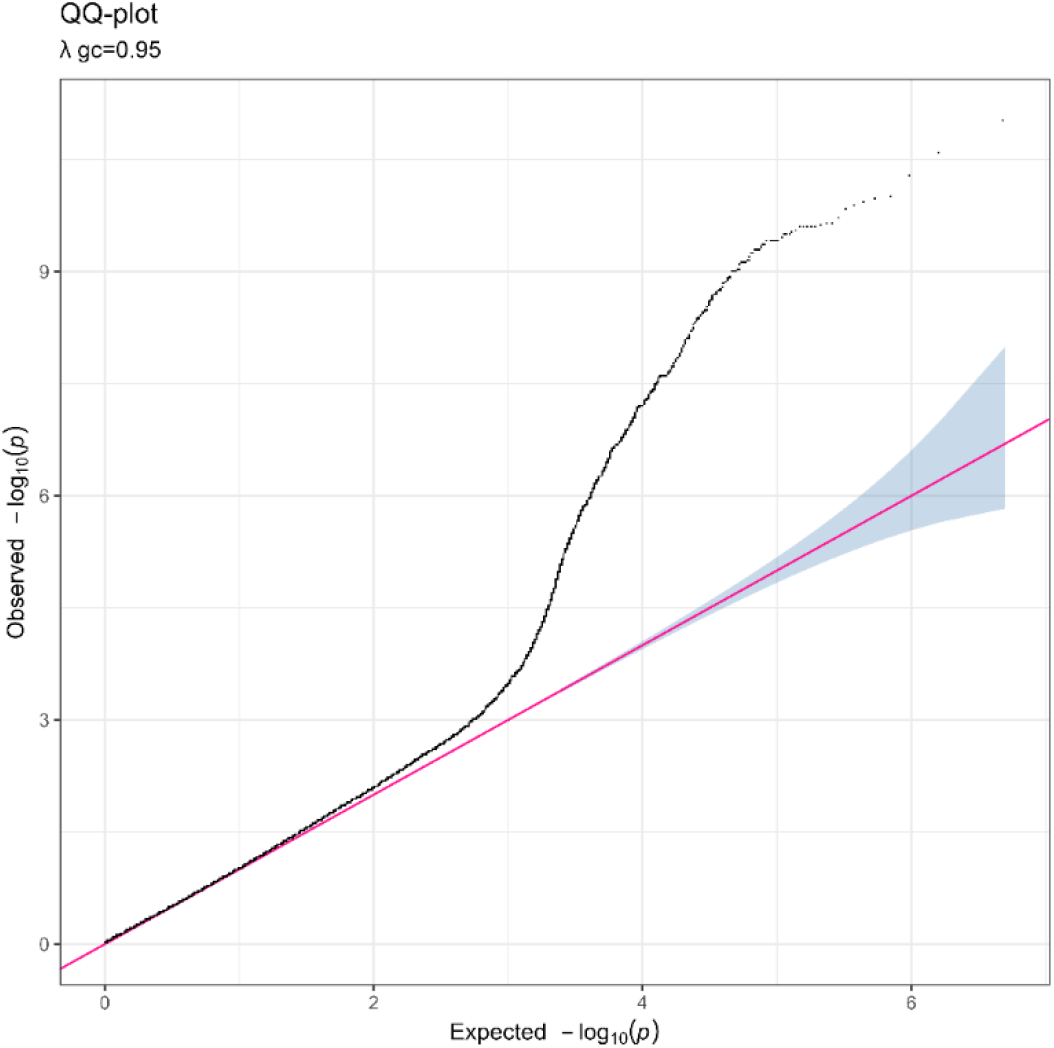
Quantile-Quantile plot for Green: Red ratio on the fourth leaf of 184 lettuce cultivars.

**Figure S10:**
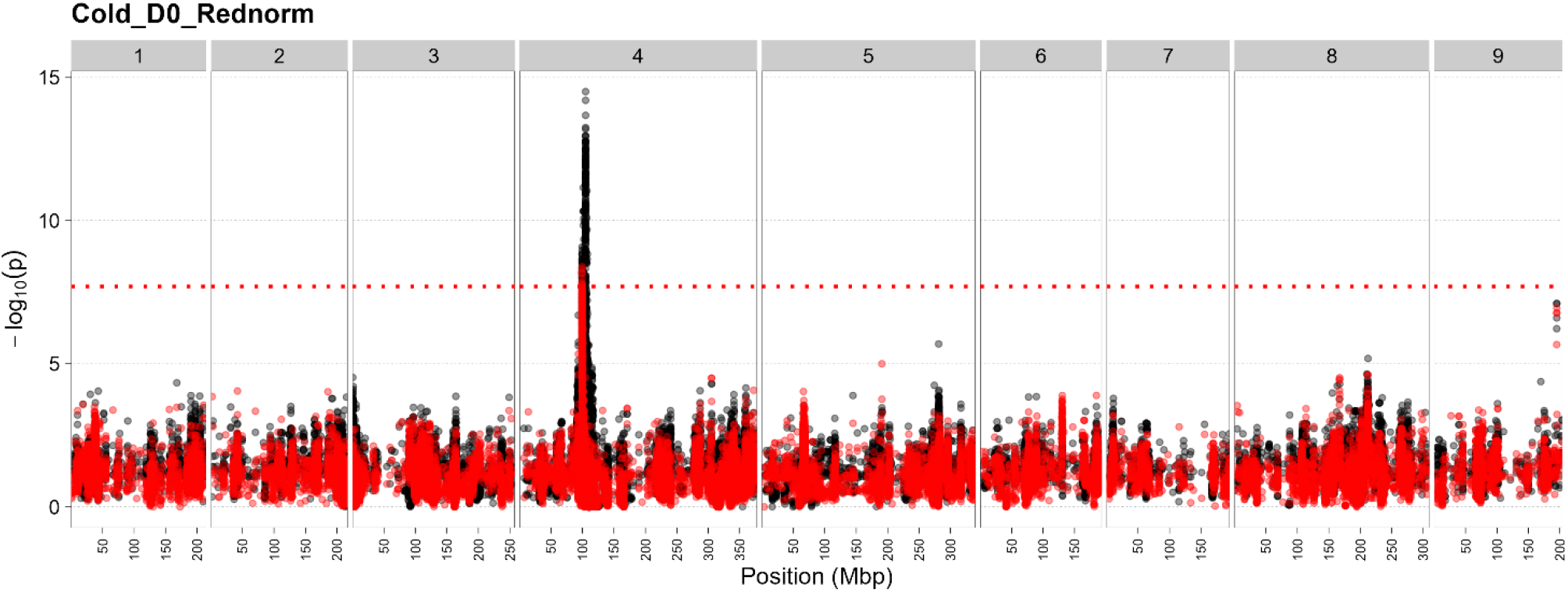
Manhattan plot of the GWAS on the Normalized Red value measured on the fourth leaf of 184 lettuce cultivars. Genomic position, indicated in megabasepairs (Mbp), is shown on the x-axis, chromosome numbers are indicated on top. Significance as –log_10_(p) is shown on the y-axis. The Bonferroni threshold of -log_10_(p) > 7.69 is indicated by the red horizontal line.

## Protocol S1: GreenLeafVI user guide

### Imaging setup and white balancing

The main limitation to image-based measurement is the variation in brightness and contrast between images. In order to calibrate the image brightness to reduce inter-image variation, we included an optional white-balancing step in the GreenLeafVI plugin (figure 1). The white-balancing step requires that a white reference area be photographed under the same conditions as the images to be analyzed. This ensures that all images are white balanced to the same reference, reducing the variation in brightness between images that can arise from differential lighting or exposure settings during the imaging process. For this purpose, we recommend including a small white reference square or circle of a known size (e.g. 4 cm^2^) in each of the images. This will serve as a reference for both the white balancing and for setting the image scale (via Analyze > set scale), which is useful for measuring the leaf area in addition to its Chl content. Alternatively, a single white reference area could be photographed at the start or end of the series of images to calibrate the entire set, assuming that lighting and exposure conditions are constant throughout the process.

When running the white balancing step, the user can select an input- and output folder and can select whether the images should be calibrated individually or batch-calibrated based on a single white reference image. Individual white balancing requires the user to select the reference area in each of the images and is therefore more time-consuming than the batch calibration, but is also more accurate and suitable for comparing images where the lighting, exposure, and focal distances are not constant. After the user has chosen whether to process images individually or in batch, the image is split into Red, Green, and Blue channels and the mean pixel intensity in the each channel is automatically calculated in the selected white area. Next, the pixel adjustment factor for each of the channels is calculated by dividing the maximum brightness of 8-bit images (255) by the mean pixel intensity of the white area (adj_factor = 255/mean). The white reference is then set to an intensity of 255 in all three channels, and the pixels outside of the reference area are then normalized based on the adjustment factor. Finally, the image is re-stacked to an 8-bit RGB image and saved in a folder selected by the user, and the image is saved under its original name with addition of a “_whitebalanced” suffix.

For optimal reproducibility between images, we recommend imaging under constant lighting with a uniform background that shows sufficient contrast to the leaves that will be imaged such as a black, white, light blue, or orange background. In addition, variation between images can be further reduced by using the same focal distance between the camera and the leaves or plants that will be imaged throughout the process. If multiple objects are to be measured in one image, it is important that these do not overlap if a separate measurement of each object is desired.

### Segmentation of images to reduce background

In order to exclude measurements of non-leaf objects or areas in the background of the image, we included an optional segmentation step in the GreenLeafVI plugin (figure 1). This process uses the difference in color between the leaves and the background to extract only leaf-like objects. Because of this, the segmentation performs better when a background with an even color that has sufficient contrast to the leaves is used during imaging, such as black, white, light blue, or orange.

In order to run the segmentation process, the user needs to select an input- and output folder, and can adjust the settings for minimum object area, as well as minimum- and maximum values in the hue saturation value color space (HSV). While the settings included in our plugin should perform well for any leaf-like object, we recommend that users customize these settings to the object of their interest to optimize segmentation performance. When the segmentation step is performed, a mask is created based on the HSV and minimum area inputs, and used to remove the background which is set to 0. The images are then saved in a folder selected by the user under their original name extended by a “_segmented” suffix.

### RGB analysis and color-based methods of chlorophyll estimation

The main feature of the GreenLeafVI plugin is the measurement of pixel intensity of individual objects (leaves) in the Red, Green, and Blue channels and the calculation of various colorimetric visual indexes (CVIs) from these data (figure 1C). The RedGreenBlue measurement can be run on all images saved in a single folder at once (we analyzed up to 446 images at a time), and the results are saved as a.csv file in the same folder. Images that can be used as input are preferably white-balanced and segmented, but the measurements can be run on unprocessed images as well, although this may result in poor identification of individual objects.

To measure the R, G, and B pixel intensities of individual objects in an image, a mask is made that selects individual objects in a similar manner as in the segmentation step. It then splits the original image into Red, Green, and Blue channels and overlays the mask on each of the channels. The minimum, maximum, median, and mean pixel intensity of each object of the mask is then measured for each of the channels, and written to the results file. The mean values of the Red, Green, and Blue channels are used to calculate the different visual indexes that are used to estimate Chl content: Green:Red ratio (GR_ratio), Red:Green ratio (RG_ratio), Kawashima index (Kawashima), Green Leaf Index (GLI), Normalized Difference Index (NDI), normalized Red (Red_norm), normalized Green (Green_norm), normalized Blue (Blue_norm), and the Woebbecke index (Woebbecke). The results file also contains a column with the image name, the region of interest (ROI), which refers to individual objects in the image, their area, and an x- and y-coordinate that can be used to retrace to which object a measurement refers.

Since ImageJ scans for objects in a left-to-right and top-to-bottom order, the top left object will be the first to be detected and the bottom-right the last. This can be taken into account when preparing multiple leaves in one image, for example by making sure that the objects are ordered top-to-bottom or by arranging the objects in a diagonal from left to right (figure 1). Alternatively, each leaf or plant can be imaged individually. For high-throughput purposes, we advise creating a metadata file that links the image name to the content of the image, which can then be merged with the results file for further analysis.

## Notes

### Competing Interest Statement

The authors have declared no competing interest.

### Funding

- Dutch Research Council: GSGT.2019.024, P17-19

